# A short-term decellularisation technique for porcine carotid arteries that conveys a structural stimulus to cells

**DOI:** 10.1101/2022.06.28.497956

**Authors:** E. Fitzpatrick, R. Gaul, C. Smekens, P. Mathieu, B. Meehan, B. Tornifoglio, P.A. Cahill, C. Lally

## Abstract

In many cases, treatment for stenotic atherosclerotic lesions requires the use of bypass grafts to divert blood flow around the diseased vessel sections. Autologous vessels are considered the “gold standard” for bypass conduits; however, the shortage of healthy autologous vessels has resulted in an increasing focus on optimising synthetic, biological and/or tissue engineered vascular bypass grafts. While many of the previously published methods have been shown to fall short of producing an ideal TEVG, this report presents a decellularisation process that produces an acellular vascular graft that is efficient, cost effective, and could be readily automated. The resulting graft can be used “off the shelf”, has preserved arterial structure and mechanical properties, and conforms to decellularisation criteria regarding the sufficient removal of cellular and genetic components. Additionally, the graft does not require any priming, supports molecular transport, can withstand supraphysiological pressures, and can support cell attachment and growth under physiological strain conditions whilst providing structural cues for cell adhesion and growth.

**Impact statement:** Vascular disease remains the leading cause of mortality worldwide. In the absence of suitable autologous vessels, there currently exists a clear clinical need for ‘off the shelf’ vascular grafts that can successfully bypass diseased arteries. This paper outlines a short-term method for obtaining such a graft. The technique used involves decellularising porcine carotid arteries whilst preserving arterial structure and mechanical properties and is compliant with the international standard for implantable vascular prostheses: EN ISO 7198:2017. Additionally, this protocol is cost and time effective, and produces reproducible “ready to use” acellular grafts that support molecular transport, can withstand supraphysiological pressures, and can support cell attachment and growth with controlled structural cues under physiological strain conditions.

## 1.0 Introduction

Atherosclerosis causes a reduction in the vascular luminal area (stenosis) by the aggregation of cells, cellular debris, and lipoproteins. This can result in tissue/organ ischemia and thrombosis which are observed clinically as unstable angina, critical limb ischemia, myocardial infarction, and stroke [1,2]. Currently, treatment includes bypassing severely stenosed vessels using grafts to redirect blood flow, and coronary and peripheral artery bypass grafts are commonly used to relieve symptoms of unstable angina and critical limb ischemia [1]. Given their inherent biocompatibility, autologous vessels such as the saphenous vein and internal mammary artery represent the “gold standard” in bypass graft materials. Crucially, they have shown superior patency when compared to synthetic graft materials in various vascular settings within the peripheral vasculature [3]. Unfortunately, autologous vessels are often rendered unsuitable for use due to underlying vascular disease or previous use in bypass surgeries [4]. Additionally, the surgery required to obtain autologous vessels is invasive and fraught with potential complications such as infection and pseudoaneurysms [5]. To overcome this, there are three main categories of alternative graft production currently under examination for use. Firstly, the fabrication of prosthetic grafts using synthetic and/or natural polymers. Customised fabrication techniques aim to emulate the structural organisation of the native extracellular matrix (ECM) using synthetic and/or biological materials [6, 7]. This category of graft has been in circulation since the 1950s; however, despite the significant scientific advances in all pertinent fields (including material science, pharmacology, and device fabrication), these grafts have not significantly increased patient survival rates to-date [8]. It is speculated that this may be due to their limitations related to their integration into the body, susceptibility to infections, inadequate biomechanical function and cytotoxic degradation *in vivo* [8, 9]. Furthermore, compliance mismatch between synthetic prostheses and the neighbouring vessel wall can compromise long-term patency, ultimately leading to premature graft failure [4, 9].

The second category of alternative grafts are cell/tissue self-assembled conduits, also known as a “bottom-up” tissue engineered grafts [5, 10]. While promising, “bottom-up” tissue engineered vessels can have with issues arising in their mechanical integrity stemming from cell-associated complications, such as difficulties in controlling cell-aggregation and ECM production and deposition *in vitro* [11].

Finally, the third category of alternative grafts are decellularised vessels. The decellularisation method, also known as a “top-down” tissue engineering approach, offers a method that potentially circumvents the issues associated with fabricating complex 3D architectures *in vitro* [12]. This method involves using a combination of physical, chemical and enzymatic treatments to remove cellular material from tissues, leaving behind a non-immunogenic porous ECM [13]. However, decellularisation, to-date, has also failed to yield an ideal TEVG. Issues arise in finding the balance between the effective removal of genetic and cellular remnants and maintaining arterial structure. In these cases, inadequate decellularisation results in adverse immune reactions and graft rejection, while adequately decellularised grafts can exhibit compromised arterial structure and biomechanics often resulting in graft related thrombosis and aneurysm [12, 14-17,63].

This paper presents a decellularisation protocol that overcomes these issues, producing a graft with maintained structural integrity, appropriate mechanical properties, and compliance with decellularisation criteria (< 50ng dsDNA/mg dry weight of tissue and removal of cellular content as examined by histochemical analysis) [18, 19]. Subsequent water permeability tests conducted in accordance with ISO 7198:2016/2017 confirm that the protocol results in a graft with sufficient molecular transport without a high risk of haemorrhage and vessel inflation tests confirm it can withstand supraphysiological conditions (≤ 2,585.75 mmHg/ 50 Psi), minimising graft rupture risk. Additionally, unlike other grafts, this process produces a graft that does not require any further processing to support cell adherence and cells remain attached to tubular vessels during long term cell culture (11 days) and remain attached while subjected to physiological cyclic strain (2-8% uniaxial). This protocol is also economically attractive due to its automatable nature, cost effective chemicals, and there is a quick turnaround. Taken together, this means that ready to use grafts could be produced at minimal cost in a time effective manner to supply constant and increasing demands.

## 2.0 Methods and materials

### 2.1 Tissue isolation and cryopreservation

Fresh porcine necks from 6-month-old pigs weighing approximately 100 kg were obtained from a local slaughterhouse within 1 hour of slaughter and transported on ice to the laboratory. The common carotid arteries, approximately 4 - 6 cm in length, were excised using standard dissection tools and subsequently rinsed in PBS at room temperature (Supplementary figure 1). The vessels were either used immediately (fresh) or cryopreserved for later use. The samples were frozen at a controlled rate of −1°C/min to −80°C in a cryoprotectant medium previously described for vascular tissue consisting of 1.8 M dimethyl sulfoxide (DMSO), 0.1 M sucrose, and Roswell Park Memorial Institute 60 (RPMI-60) medium [20]. This cryopreservation method became the adopted routine tissue freezing method throughout the study. Prior to use, cryopreserved samples were thawed quickly at 37 °C in a water bath [21].

### 2.2 Decellularisation and DNAse treatment of carotid arteries

Porcine carotid arteries (PCA) were decellularised based on a previously described method for porcine coronary arteries which was adapted for carotid arteries [22]. Briefly, sections of carotid arteries approximately 4 cm in length were attached to a perfusion system using nylon barbs which were inserted into the arterial lumen (Supplementary figure 1). A 0.1 M sodium hydroxide (NaOH) solution was perfused through the lumen of the fresh and cryopreserved carotid arteries for 15 hours, while the outer adventitial layer was exposed to the NaOH by static submersion in a bath. This was followed by a 31-hour rinse in a 0.9% saline solution which was changed three times to remove any cellular debris and chemical residue. To adequately remove DNA fragments, vessels were removed from the perfused system and incubated in 3 Kunitz/ml protease-free DNAse (Worthington, LS006343) in 1x enzyme reaction buffer (40 mM Tris-HCl pH 8.0, 10 mM MgSO4, 1 mM CaCl2, 100 μg/ml primocin) for 19 hrs at 37°C. Sections from each vessel were taken after each treatment step and prepared for histological analysis. For samples that were used for confocal, the decellularised PCA were cut longitudinally in order to produce strips and the intima was removed under a dissection microscope.

### 2.3 Histological processing

Standard histological techniques were used to verify the absence of nuclear material in decellularised tissue and to investigate the effect of decellularisation on collagen, elastin, and glycosaminoglycan (GAG) structure. Briefly, sections of arteries were fixed in 4% paraformaldehyde (PFA) overnight and stored in 70% ethanol. The samples were then dehydrated using increasing concentrations of ethanol, cleared in xylene for 4 minutes, embedded in paraffin, sectioned into 7 μm thick slices, and mounted onto slides. Regardless of histological stain used, all samples were subjected to paraffin rehydration prior to staining. This process was conducted at room temperature and involved immersing the slides in xylene for 4 min, followed by submersion in 100% ethanol for 4 min and subsequently deionized water for 1 min. Staining was then immediately conducted in singular batches using a Leica 5010 autostainer. Sections were stained with Haematoxylin and Eosin (H&E), Verhoeff’s elastin stain, picrosirius red, and Alcian blue prior to being permanently mounted in DPX. Brightfield imaging was done using an Olympus EX-41 microscope. Two polarised light microscopy images of the picrosirius red stained samples were merged using an in-house MATLAB program. Slices were also stained with 4’, 6-diamidino-2-phenylindole (DAPI) containing mounting media (Fluoroshield F6057, Sigma) to visualise nuclear material and were imaged using an Olympus IX-83 fluorescent microscope equipped with an Olympus DP50 camera. Slides were then permanently sealed using clear nail varnish.

### 2.4 Collagen alignment

Sample sections designated for collagen staining were incubated in pre-warmed media (GIBCO DMEM + GlutaMAX basal media, 10% FBS and 100 μg/ml primocin) supplemented with 100 μg/ml CNA35-488/546 for 2 hrs at 37 °C. Samples were then washed twice in 1x PBS before being fixed at 4°C overnight in 10% formalin. Following this, samples were washed in 1x PBS and stored at 4°C until imaged using a Leica SP8 confocal microscope. Each vessel was imaged in ≥ 4 locations. Images were then processed using the OrientationJ Distribution plugin in ImageJ, the distribution of orientation was normalised, entered into GraphPad Prism version 6 and plotted for a visual representation of the collagen orientation.

### 2.5 Collagen content

Collagen levels were quantified by measuring the amount of hydroxyproline in fresh, decellularised and DNAse treated decellularised samples. Briefly, papain-digested samples were further digested in 38% HCl at 110°C overnight. The samples were then cooled to room temperature before being centrifuged at 15,000 x G for 10 mins at room temperature. In a fume hood, sample lids were removed, and samples were then placed in a heating block set to 50°C until all the liquid had evaporated off. Samples were dissolved in ultrapure H_2_O, vortexed and centrifuged using a mini desktop centrifuge. Samples were either assessed immediately or stored at –20°C until use. The hydroxyproline assay was then conducted in accordance with manufacturers recommendations (Sigma, MAK008), and the reagents were used within 2 – 3 hours of preparation. Briefly, samples and standards were incubated in a 96 well plate with oxidation buffer and chloramine-T reagent for 20 mins at room temperature and protected from light. DMBA (dimethylaminobenzaldehyde) reagent was then added and pipetted until the solution became clear. The plate was then sealed (Sigma, Z369667) and incubated at 60°C for 20 mins. Hydroxyproline content was measured at a wavelength of 570 nm and the amount of collagen present in each sample was calculated using the known value of 13% hydroxyproline in collagen [23]. The average and standard deviation were entered into GraphPad Prism version 6 and plotted for a visual representation of the data. One-way ANOVA statistical analysis was subsequently conducted on the dataset.

### 2.6 DNA quantification

Samples for DNA content quantification were flash frozen and stored at −20°C until ready for freeze drying. Samples were introduced into a precooled (−55°C) freeze drier for 1 hour. The temperature was then increased by 1°C/min to −40°C, at which point it was held for 1 hour and a vacuum (200 mTorr) was applied. The temperature increased 1°C/6 min until it reached 0°C and then held for a further 17 hours. The temperature was increased 1°C/min to 26°C under vacuum and held until ready to be removed. Samples were papain digested using 125 μg/ml papain in activated papain digestion buffer (0.05 M EDTA, 0.1 M sodium acetate, 5 mM L-cysteine-HCl, pH 6.0) for 18 hours at 60°C, 10 RPM. Digests were vortexed and centrifuged at 15,000 x G for 10 minutes at 4°C prior to immediate DNA quantification. DNA quantification analysis was conducted using a Quant-iT^™^ Picogreen^™^ dsDNA quantification kit (Life Technologies) in accordance with the manufacturers protocol. Briefly, papain-digested samples and standards were incubated with picogreen reagent for 5 minutes at room temperature and the fluorescence measured at an excitation wavelength of 485 nm and emission wavelength 530 nm. DNA concentrations were then extrapolated from a standard curve and the average and standard deviation of the readings were obtained and entered GraphPad Prism version 6. The data was then plotted and analysed using a two-tailed (unpaired) t-test (decellularised vs. decellularised + DNAse treated).

### 2.7 Water permeability and pressurisation

The water permeability was determined in accordance with methods outlined in the industrial and international standard: ISO 7198:2016/2017. Water permeability assessment was carried out with grafts attached to a static pressure head (positioned to achieve 120 mmHg). The end of the pressurised system was closed off and any water that permeated through the graft was collected in a closed container. The sealed container was replaced every 30 mins for 2 hrs and the permeated water was measured with the final 3 readings averaged and inserted into the graft permeability equation, P=Q/T, where P is the graft permeability [ml/cm^2^/min], Q is the permeated fluid volume per area of graft [ml/cm^2^], and T is the time of collection [min]. The area of the graft [cm^2^] was calculated, A=D*L\**π*, where D was the inner diameter of the graft [cm] and L the length of the graft [cm]. Grafts were then subjected to the supraphysiological pressure of 50 psi (2,585.75 mmHg) using a MERITMEDICAL BasixCompak^™^ Inflation Device.

### 2.8 Mechanical Testing

Common carotid arteries (n=7) were divided into 3 proximal and 3 distal ring sections of approximately 2mm axial length, shown schematically in Supplementary figure 2. The outer 2 mm vessel sections, on both proximal and distal sides, were subjected to a uniaxial tensile test on the day that the vessel was taken out of cryopreservation. For every vessel, these outer sections served as controls (denoted by “C”). The remaining part of the vessels was decellularised and further segmented to obtain both the proximal and distal decellularised segments. These were tested immediately after the decellularisation process was finished (sections denoted by “D”). The remaining vessel sections were treated with the DNAse reaction buffer, segmented to get proximal and distal rings, and mechanically tested (sections denoted by “DD”). Uniaxial tensile tests were conducted similar to the method described by Sheridan et al. [24] using a Zwick tensile testing machine (Zwick Z005, Roell, Germany) with a 100 N load cell. Samples, 2 mm in length, were placed on purpose-built grips such that both 1 mm diameter pins projected through the arterial lumen, as shown in Supplementary figure 3. Once placed on the grips, the force was zeroed. Like previous studies [24, 25], samples were tested in the circumferential direction through vertical displacement of the crosshead at a constant rate of 60%/min. A preload of 0.05 N was applied to remove rigid body motion from the ring test before 5 preconditioning cycles up to 10% grip-to-grip separation were applied to minimise viscous effects from the vessel. Samples were tested to failure at a strain rate of 60%/min of the recalculated initial gauge length. The force-extension data recorded by the Zwick machine was then converted to true stress, *σ* and true strain, *ε* values in MATLAB (Mathworks) using Equation 1 and 2 below.

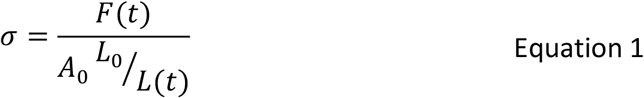

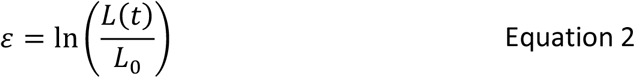

where F(t) is the measured force at time t, *A*_o_ is the initial cross-sectional area (CSA) of the ring sample, *L*_o_ is the initial gauge length after the preload and *L*(*t*) is the gauge length at time t. The CSA was determined as twice the undeformed specimen wall thickness multiplied by the sample width. The associated incremental moduli of the elastin and collagen dominant phases were calculated as the slope of the stress-strain curve in the initial and final regions of the curve, respectively (Supplementary figure 3). Moreover, the tissue distensibility was evaluated by extrapolation of the linear trendline in the high-strain region to intercept the x-axis. The ultimate tensile strength (UTS) was also recorded for each sample, where the UTS was defined as the point at which the observed stress measurements began to decrease. This data was then graphed using MATLAB.

During statistical analysis, both proximal and distal results of each vessel condition were treated separately. The data is presented as mean ± standard deviation with n=7 for each vessel condition (C, D, and DD) and vessel side (proximal or distal). Outlier detection was performed (OD) by means of GraphPad Prism’s build-in ROUT outlier detection method. This resulted in the removal of 1 set (C, D and DD) of distal vessel sections (vessel n°3). Tests on vessel sections of the same vessel were regarded as repeated measurements. Samples from different arteries were considered independent. As such, for repeated measurements two-way ANOVA tests were performed in GraphPad Prism 6 (*p<0.05 considered significant), followed by Tukey’s multiple comparisons tests.

### 2.9 Isolation and culture of pbMSC

Porcine bone marrow derived mesenchymal stem cells (pbMSC) were isolated from fresh porcine femoral bone marrow. The femoral bone marrow was extracted from the femoral shaft and transferred into pre-warmed MSC expansion media (GIBCO DMEM + GlutaMAX basal media, 10 % FBS and 2 % penicillin streptomycin). Following trituration using a 16-gauge needle, the cell suspension was centrifuged at 650 g for 5 minutes and the supernatant discarded. The cell pellet was resuspended in more pre-warmed expansion media and the centrifugation and supernatant discarding step repeated. The cell pellet was then resuspended in a small volume of expansion media and triturated as before and the cell suspension was then passed through a 40 μm nylon mesh. The resulting cell suspension was then treated with lymphoprep (Axis-Shield catalogue no. 1114545) in accordance with the manufacturers’ protocol in order to isolate mononuclear cells. The mononuclear cells were plated at 2.5 × 10^6^ cells/10cm in expansion media and cultured at 37°C and 5% CO^2^. After 72 hours, the cells were washed twice with 1x PBS in order to remove non-adherent cells and debris. Flasks were seeded at 5×10^3^ cells/cm^2^ and maintained at a low density. All experiments were conducted using the first 4 cell passages.

### 2.10 *In vitro* cell seeding of planar strips

Cryopreserved longitudinally cut open strips (decellularised + DNAse treated) with the intima removed were defrosted in a 37°C water bath and washed twice with 1x PBS before being placed in 100% ethanol for a minimum of 1 hour for sterilisation. The decellularised PCA strips were then introduced into a Class II biological safety cabinet and washed twice in sterile 1x PBS before being pinned flat onto UV sterilised PDMS coated culture dishes using metal pins sterilised in 100% ethanol. A suspension of pbMSC was then seeded directly onto the strips and after 2 hours the strips were relocated into a culture dish and cultured for 48 hours in 2-10 ng/ml TGFβ1 and 100 μg/ml primocin supplemented expansion media. Following this, the PCA strips were then relocated into a BOSE ElectroForce BioDynamic 5270 bioreactor and exposed to cyclic strain of 2–8 % at 1 Hz for 72 hours. During the 72 hour strain timeframe, the cells were cultured in expansion media supplemented with 100 μg/ml primocin (without TGFβ1).

### 2.11 *In vitro* cell seeding of tubular scaffolds

Tubular vessels (decellularised + DNAse treated) were turned inside out, the intima removed and re-inverted before being stored in tissue freezing media at −20°C until ready for use. The tubular PCA were then defrosted in a 37°C water bath and washed twice with 1x PBS before being placed in 100% ethanol for a minimum of 1 hour for sterilisation. The scaffolds were then introduced into a Class II biological safety cabinet and washed twice in sterile 1x PBS. Vessels of identical lengths and diameters were incubated in a suspension of 10×10^6^ pbMSC/25 ml expansion media while rotating slowly for 3 days prior to a 1x PBS wash, and cultured for 11 days with media changes every 3-4 days.

### 2.12 Immunocytochemistry

Following culture, the recellularised PCA were washed in 1x PBS before being stained for collagen by incubation in pre-warmed expansion media supplemented with CNA35-546 for 2 hours at 37°C. The samples were then washed twice in 1x PBS before being fixed at 4 °C overnight in 10% formalin. Following this the samples were washed in 1x PBS and permeabilised with 0.1% Triton X-100 PBS for 10 mins and washed twice in 1x PBS. The samples were then blocked using 5% BSA PBS at room temperature for 1 hour before being washed twice in 1x PBS. Samples were then incubated in rabbit anti-SM22α (ab14106) for 2 hours at room temperature before being washed twice with 1x PBS. The samples were then incubated in anti-rabbit alexa fluor 488 (A11008) for 2 hours at room temperature before being washed twice with 1x PBS and being stored in 1:5000 DAPI PBS at 4°C until imaged using a Leica SP8 confocal microscope. Images were then processed using Leica X software.

## 3.0 Results

### 3.1 Compliance with decellularisation requirements

Histological examination of treated vessels led to several findings. Firstly, the decellularisation technique adequately removes cellular content without noticeable damage to the underlying vascular ECM. Both H&E, see Figure 1, and Verhoeff’s staining, see Figure 2, show that following decellularisation with and without DNAse treatment, the underlying microstructure remains intact without any apparent damage. Additionally, Alcian blue staining was conducted to visually assess whether the process impacted the vascular GAG profile. Figure 3 confirms there is no observable impact to the vascular GAG at any point. It was also noted that PCA sections proximal to the aortic arch had an increased elastin:collagen ratio, whereas the reverse was evident in the more distal sections (Figure 2). The decellularisation process did not affect the overall microstructural at either of these locations. H&E analysis also shows that regardless of whether the tissue was used fresh or cryopreserved prior to decellularisation, without DNAse treatment there is a visually detectable level of residual genetic material, a fact that is corroborated by DAPI visualisation and DNA quantification (Figure 1 and Figure 4). Regarding removal of residual genetic material, the efficiency of the DNAse treatment on cryopreserved decellularised vessels was assessed considering that one of the criteria for decellularised grafts is that there must be less than 50ng dsDNA/mg of tissue (dry weight) [18, 19]. DNA quantification showed that DNAse treatment significantly (p<0.0001) reduced the DNA content to within the acceptable range, with the average decellularised control reading 620.3 ng DNA/mg of tissue (dry weight, n=5) and the average DNAse treated samples reading 38.75 ng DNA/mg of tissue (dry weight, n=8) (Figure 4B). This result supports observations made in the DAPI and H&E investigation that residual genetic material is observed following decellularisation but not following DNAse treatment (Figure 1 and Figure 4). Polarised light microscopy of picrosirius red stained slices provided additional insight into the overall collagen architecture throughout the vascular layers, see Figure 5. Low magnification assessment shows no obvious damage to collagen (Figure 5) and elastin fibre (Figure 2) architecture regardless of the approach. Collagen content quantification by the hydroxyproline assay also confirmed that the protocol had no significant effect (p=0.897) on collagen levels, with the average non decellularised control reading 39.66% collagen (n=4), decellularised control reading 45.70% collagen (n=5) and the average DNAse treated decellularised samples reading 51.3% collagen (n=8) (Figure 5B). Confirmation of collagen structure present in the tunica media from a luminal viewpoint in the PCAs was also examined using CNA35, see Figure 6 [26]. CNA35 specificity and resolution made it the ideal choice for interpreting any impact on collagen structure and alignment [26]. Examination of cryopreserved decellularised vessels with and without DNAse treatment shows that collagen is present and predominantly aligned in the circumferential direction, as would be expected (Figure 6).

**Figure 1.**
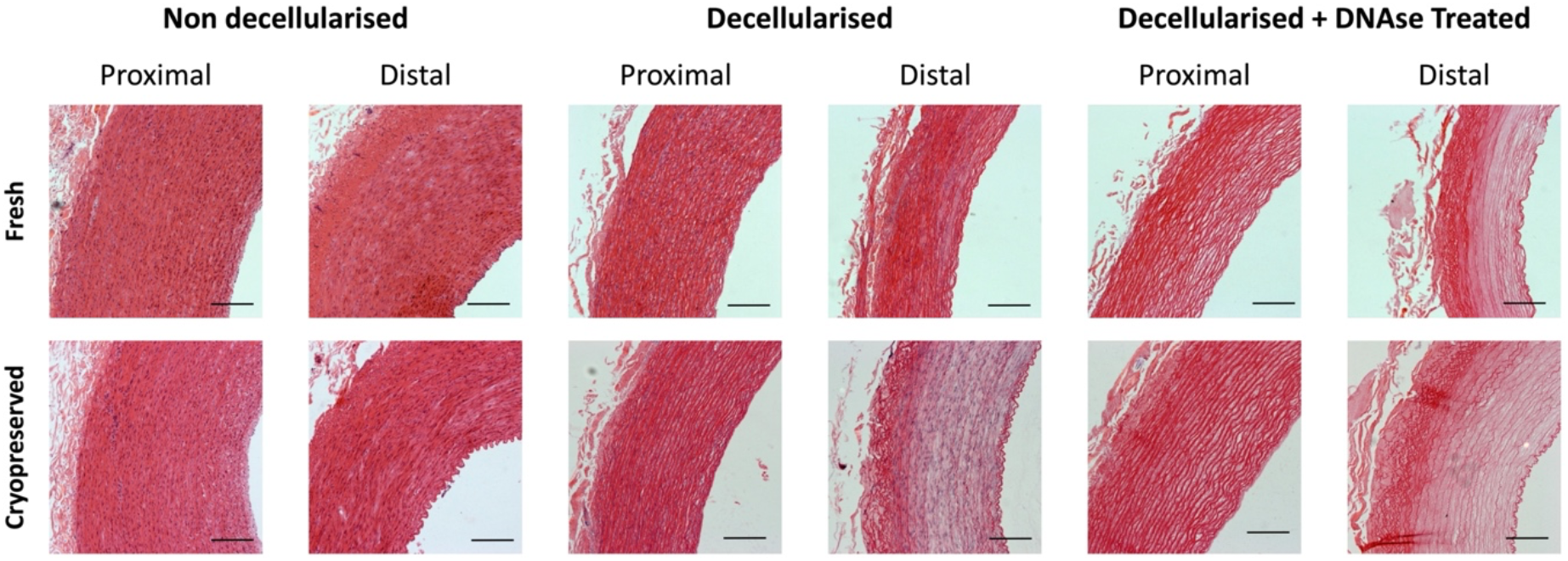
The impact of the decellularisation process on cellular content. H&E staining, scale bars = 200 *μ*m.

**Figure 2.**
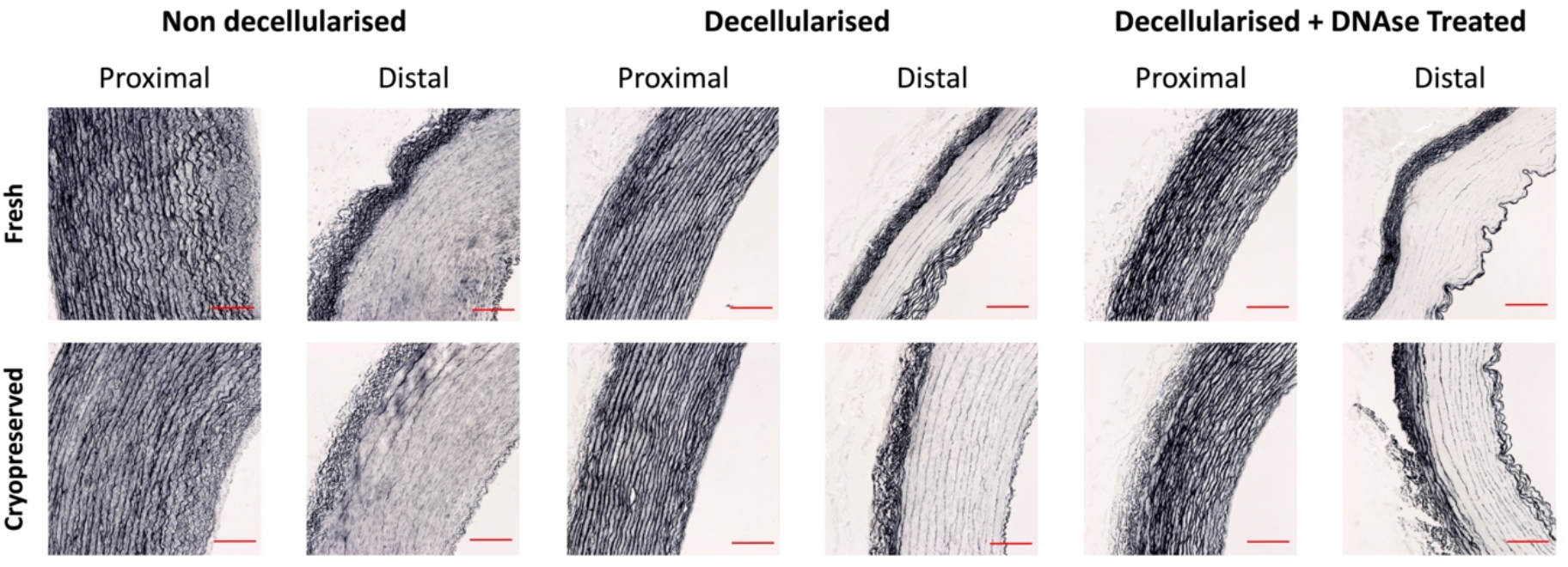
The impact of the decellularisation process on native elastin. Verhoeff’s elastin stain, scale bars = 200 *μ*m.

**Figure 3.**
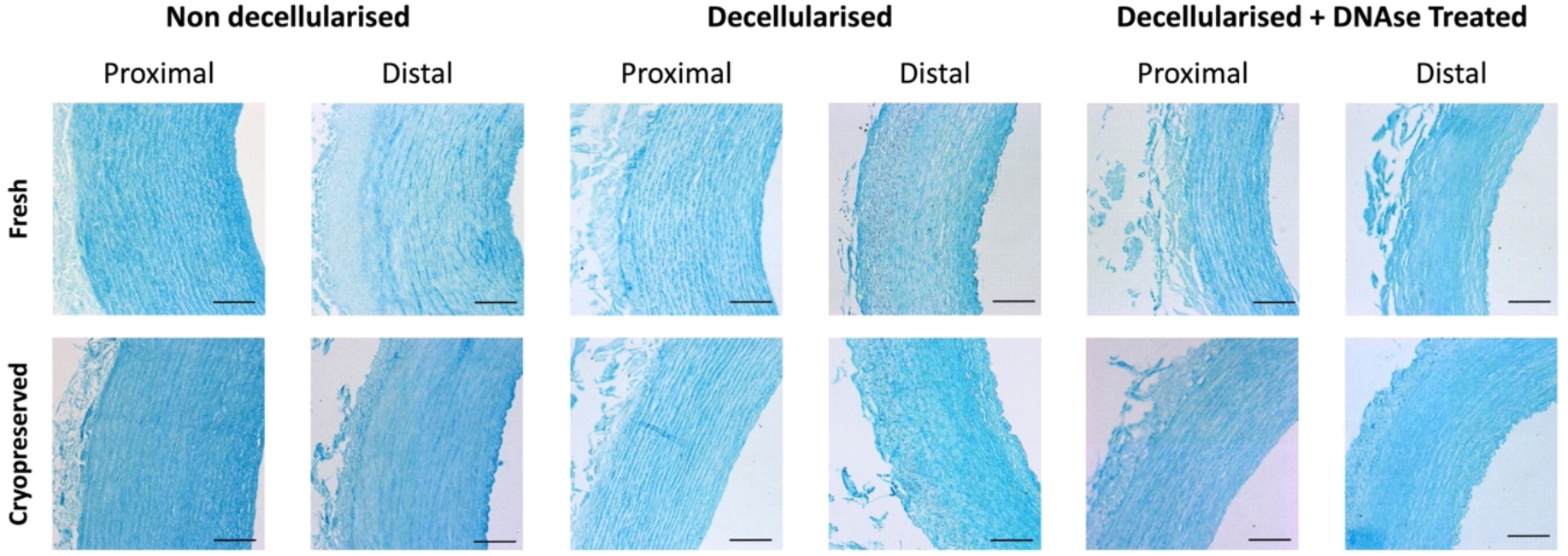
The impact of the decellularisation process on native GAG. Alcian blue stain, scale bars = 200 *μ*m.

**Figure 4.**
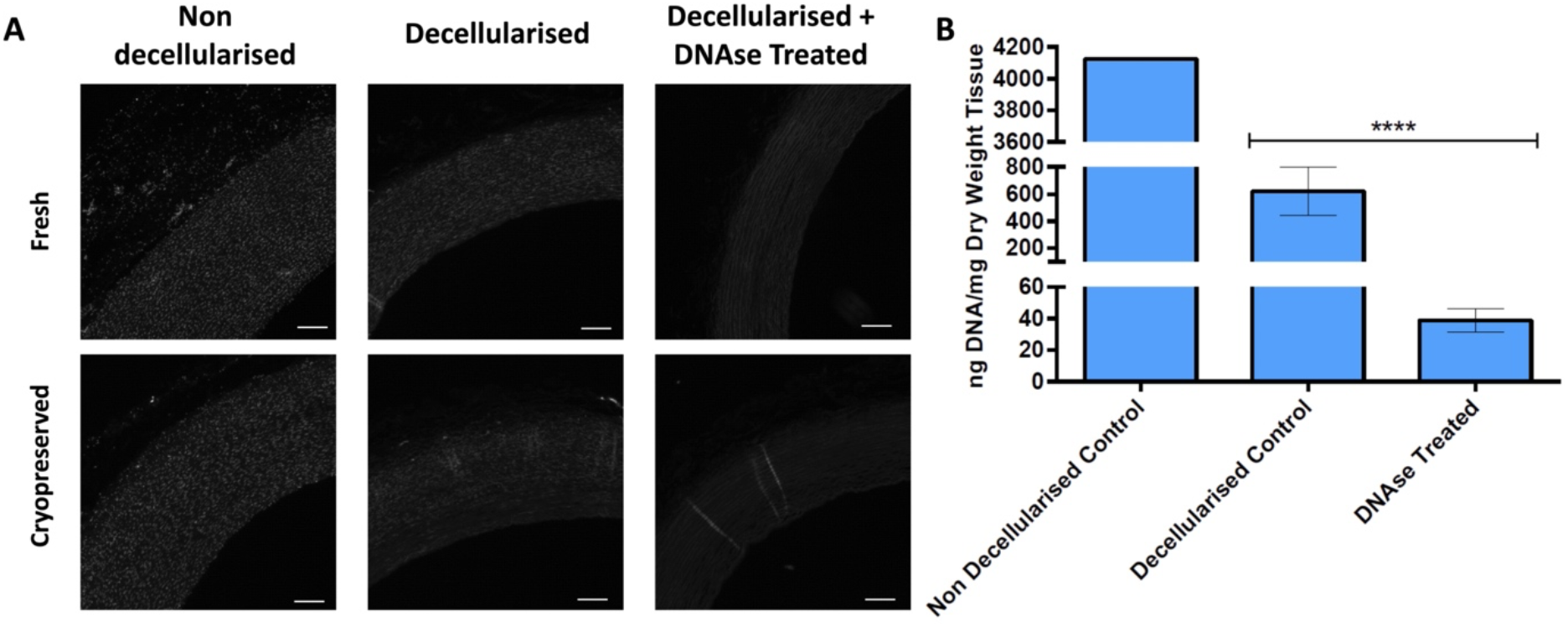
The impact of the decellularisation process on DNA content. (A) Representative axial cross-sectional fluorescent images of DNA content visualised by DAPI staining. Scale bars = 200 *μ*m. (B) DNA quantification of cryopreserved vessels showed that DNAse treatment after decellularisation yielded significant (****p<0.0001) reduction of DNA content, with a mean of 38.75 ng DNA/mg dry weight tissue (n=8) compared to decellularised only vessels (n=5) (mean of 620.3 ng DNA/ mg dry weight tissue).

**Figure 5.**
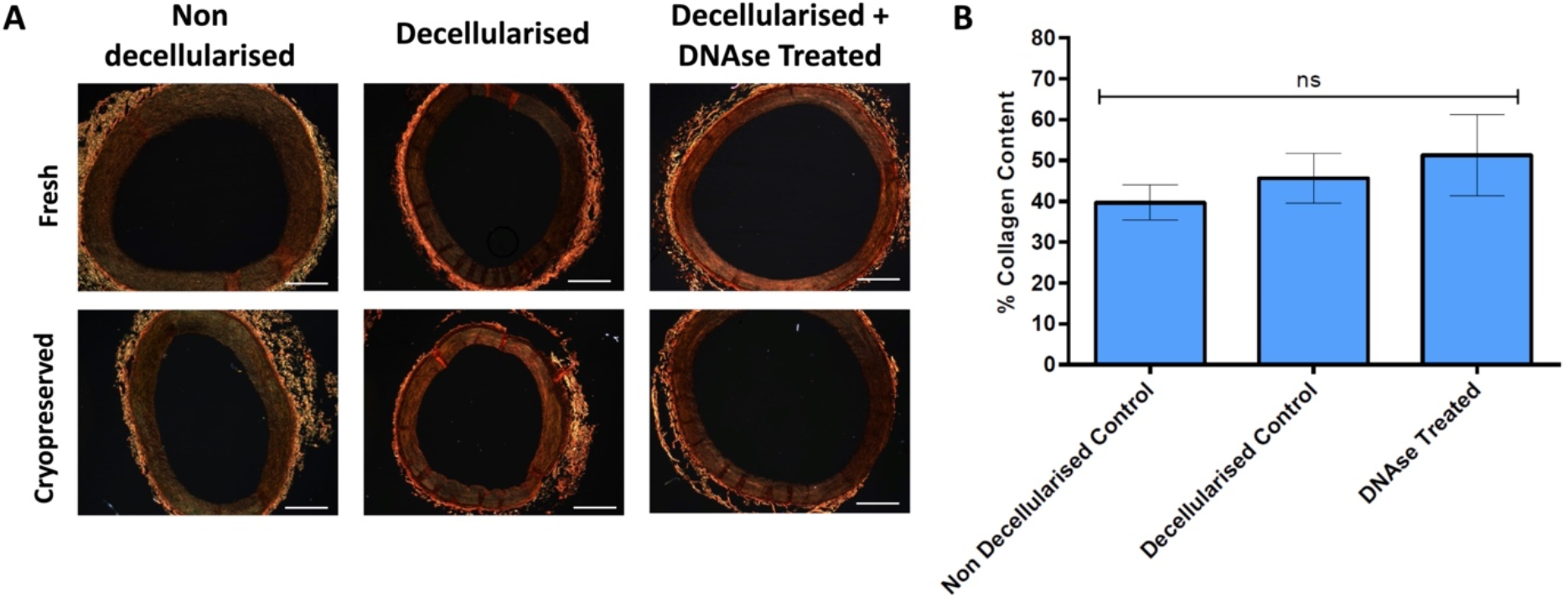
The impact of the decellularisation process on native collagen. (A) Representative low magnification axial slices of polarised light microscopy of picrosirius red stained slices. Scale bars = 500 *μ*m. (B) Collagen quantification of cryopreserved vessels throughout the protocol showed no significant effect (p=0.897) on collagen levels, with non-decellularised controls having 39.66% (n=4), decellularised having 45.7% (n=5), and DNAse treated having 51.3% (n=8) collagen content.

**Figure 6.**
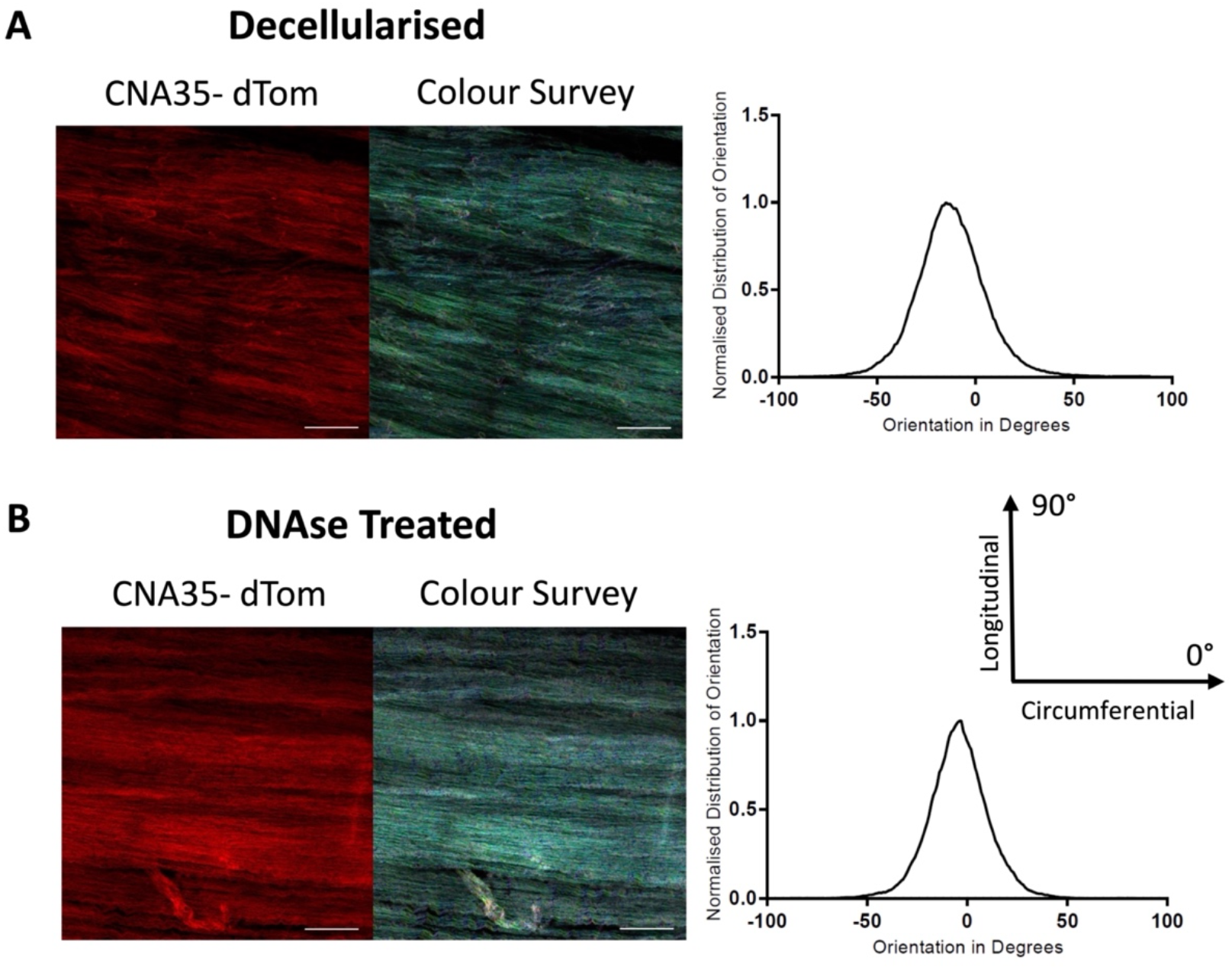
Microscopic assessment of collagen structure following decellularisation and decellularisation + DNAse treatment. Representative confocal images of the tunica media from a luminal viewpoint and the fibre direction analysis of collagen in (A) decellularised and (B) decellularised + DNAse treated cryopreserved vessels. Scale bars = 100 *μ*m.

### 3.2 Biomechanical characterisation

Water permeability characterisation is crucial for any potential in vivo graft. The rate of water permeability across the scaffolds examined was 0.009 ± 0.006 ml/cm^2^/min^-1^ (Figure 7). Jonas et al. state that porosity must be controlled to less than 50 ml/cm^2^/min^-1^ to avoid massive haemorrhage, therefore it is arguable that this graft will not haemorrhage [27]. Additionally, pressurised tests conducted on all scaffolds following the water permeability testing showed that the grafts did not fail (burst) when subjected to increasing pressure up to and including 50 psi (2,585.75 mmHg); therefore, there is no concern of potential graft failure in vivo.

**Figure 7.**
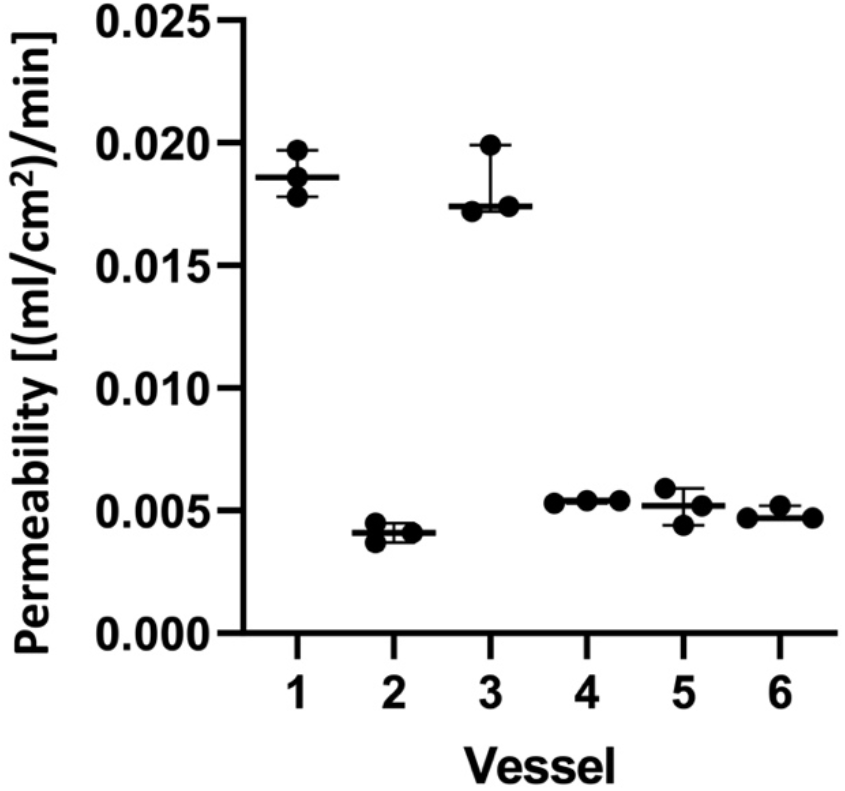
Water permeability characterisation. N=6 decellularised + DNAse treated vessels with n=3 measurements of water permeability.

Mechanically, all PCA samples exhibited a characteristic non-linear J-shaped stress-strain curve found for collagenous tissue, comprising of an early toe, transitional and linear high strain region before failure, see Figure 8A-G. However, notable variations were observed in the tissue response between native, cryopreserved-decellularised arteries, and decellularised, DNAse-treated samples (Figure 8). Additionally, differences were observed between tissue samples obtained from the proximal and distal end of the PCA. All 7 control vessels (cryopreserved, non-decellularised) showed a lengthened toe region in the proximal sections in comparison to the distal sections. This difference in the toe region was decreased in decellularised and decellularised + DNAse-treated samples and in all but one case the proximal section had an equal or greater length of the toe region. This is consistent with the increased elastin observed histologically in proximal tissue samples (Figure 2). Overall, decellularisation and DNAse treatment was not found to significantly change the tissues’ mechanical properties (Figure 9). The elastic modulus in the elastin dominated regions showed no significant differences between different treatments regardless of sample location (Figure 9A). The tissue stiffness in the collagen dominant region did increase upon decellularisation in proximal samples; however, this increase was not significant in decellularised, DNAse-treated proximal samples (Figure 9B). Additionally, the tissue stiffness in the collagen dominant region in distal sections was not significantly different across any treatments (Figure 9B). The ultimate tensile strength in distal and proximal segments showed no significant differences between different treatments (Figure 9C), and the x-intercept of the collagen dominant region also showed no significant difference between treatments in both distal and proximal sections (Figure 9D).

**Figure 8.**
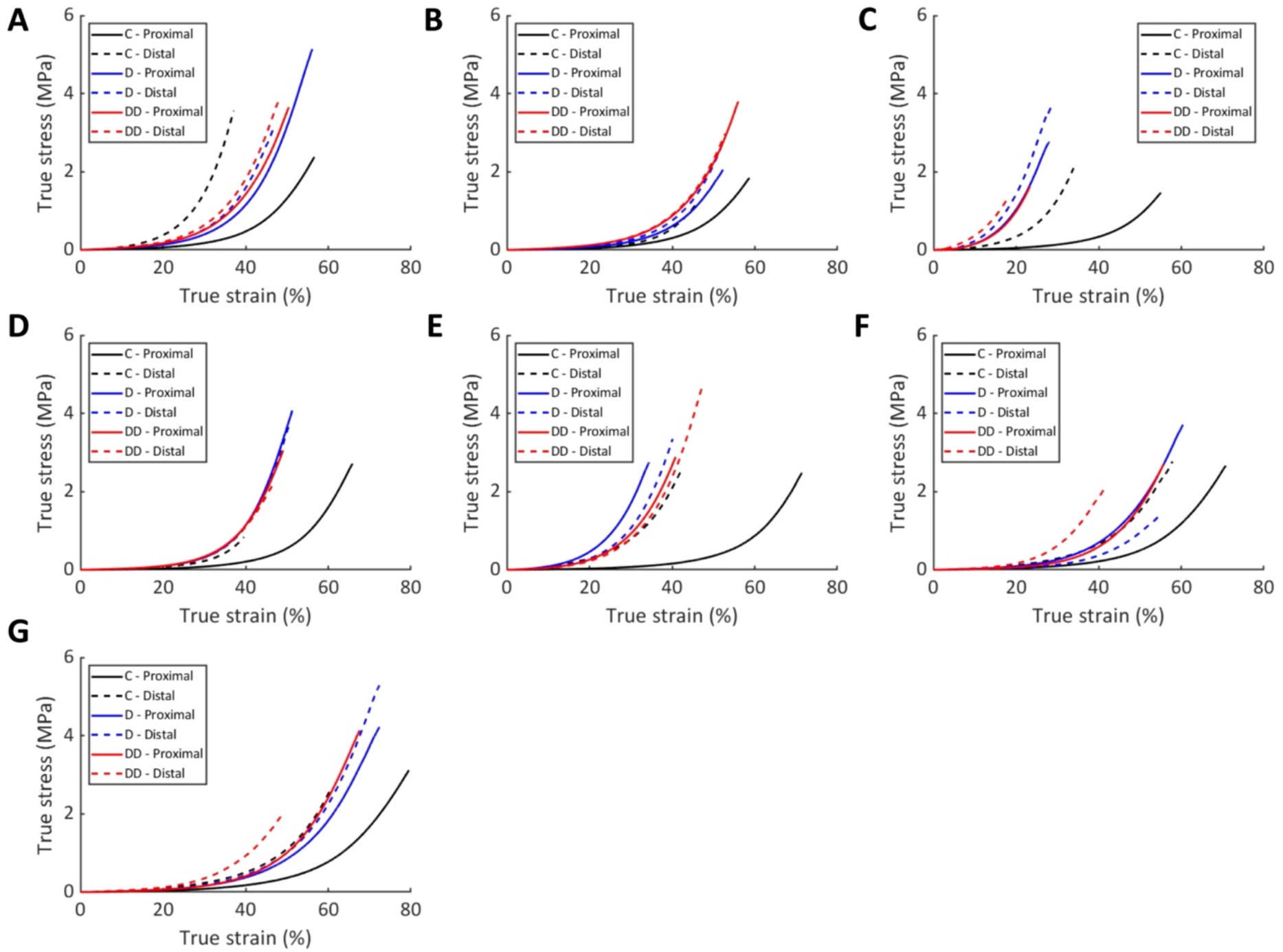
Stress-strain curves of different vessels. (A-G) Stress-strain curves for individual vessels (n=7). Cryopreserved, non-decellularised = C, cryopreserved, decellularised = D, cryopreserved, decellularised + DNAse treated = DD. Proximal and distal sections were assessed separately at each protocol condition, testing performed was uniaxial ring tensile tests up to failure.

**Figure 9.**
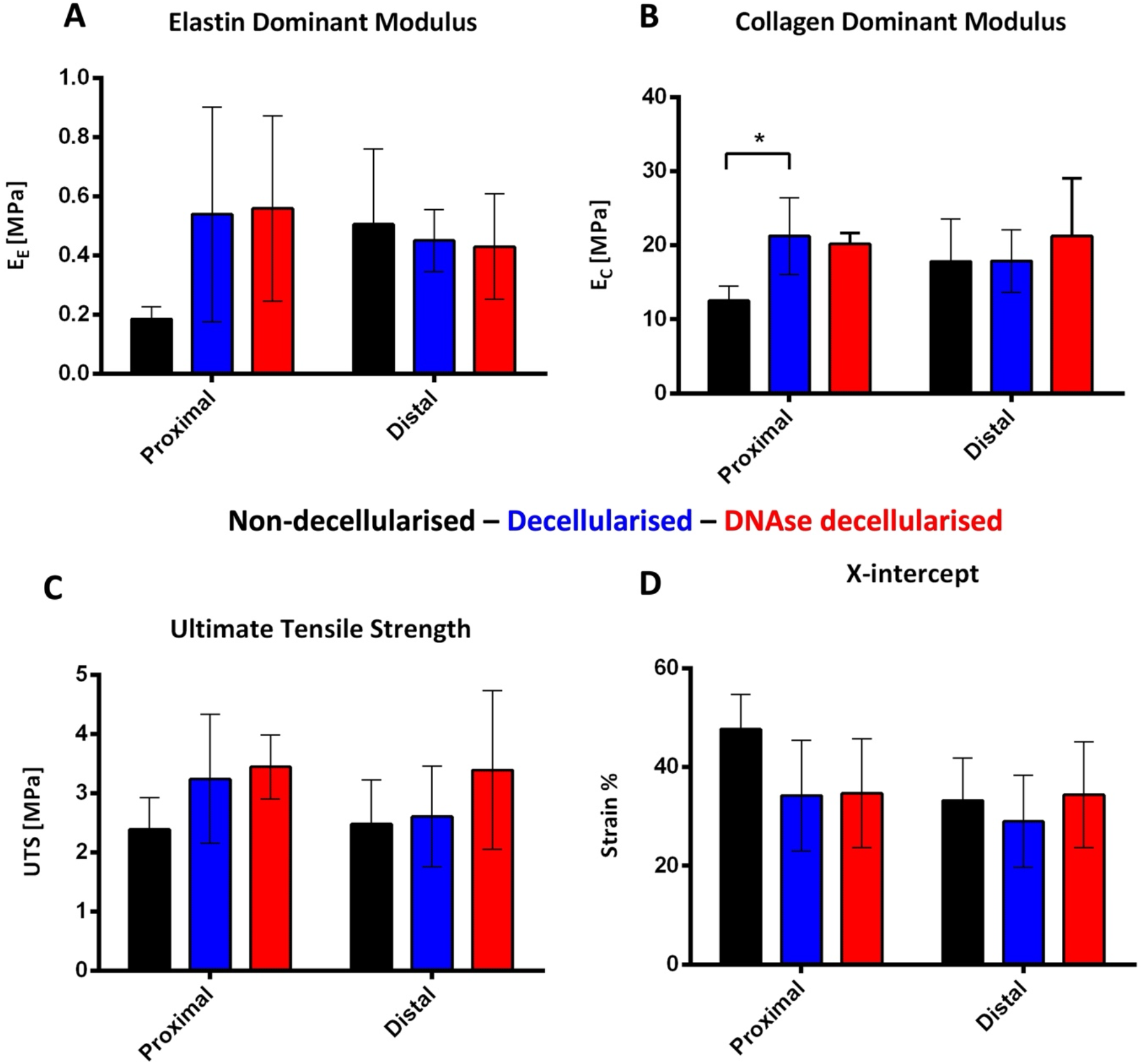
Average mechanical properties of PCAs. Mechanical properties extracted from the stress-strain curves for cryopreserved, non-decellularised (black), cryopreserved, decellularised (blue), cryopreserved, and decellularised + DNAse treated (red). (A) Elastin dominant elastic moduli, (B) collagen dominant elastic moduli, (C) ultimate tensile strength, and (D) x-intercept of collagen slope.

### 3.3 Graft repopulation

Finally, critical to potential use as a vascular graft, cells must be able to grow on the resulting scaffold. pbMSC were seeded onto planar strips of cryopreserved decellularised and DNAse treated scaffolds and cultured statically for 48 hrs in the presence of TGFβ-1 [28]. The strips were then uniaxially strained at 2–8 % for 72 hours to assess if the cells would remain on the scaffold under normal physiological strain conditions. Confocal examination shows that cells can be relatively evenly seeded onto the scaffold, see Figure 10, and that cells attach to the underlying collagen, see Figure 11. Further to this, tubular decellularised vessels were incubated in the presence of pbMSC, and confocal analysis shows that pbMSC remain adhered to the tubular constructs 11 days post seeding, confirming that the tubular vessels can also be repopulated, see Figure 12.

**Figure 10.**
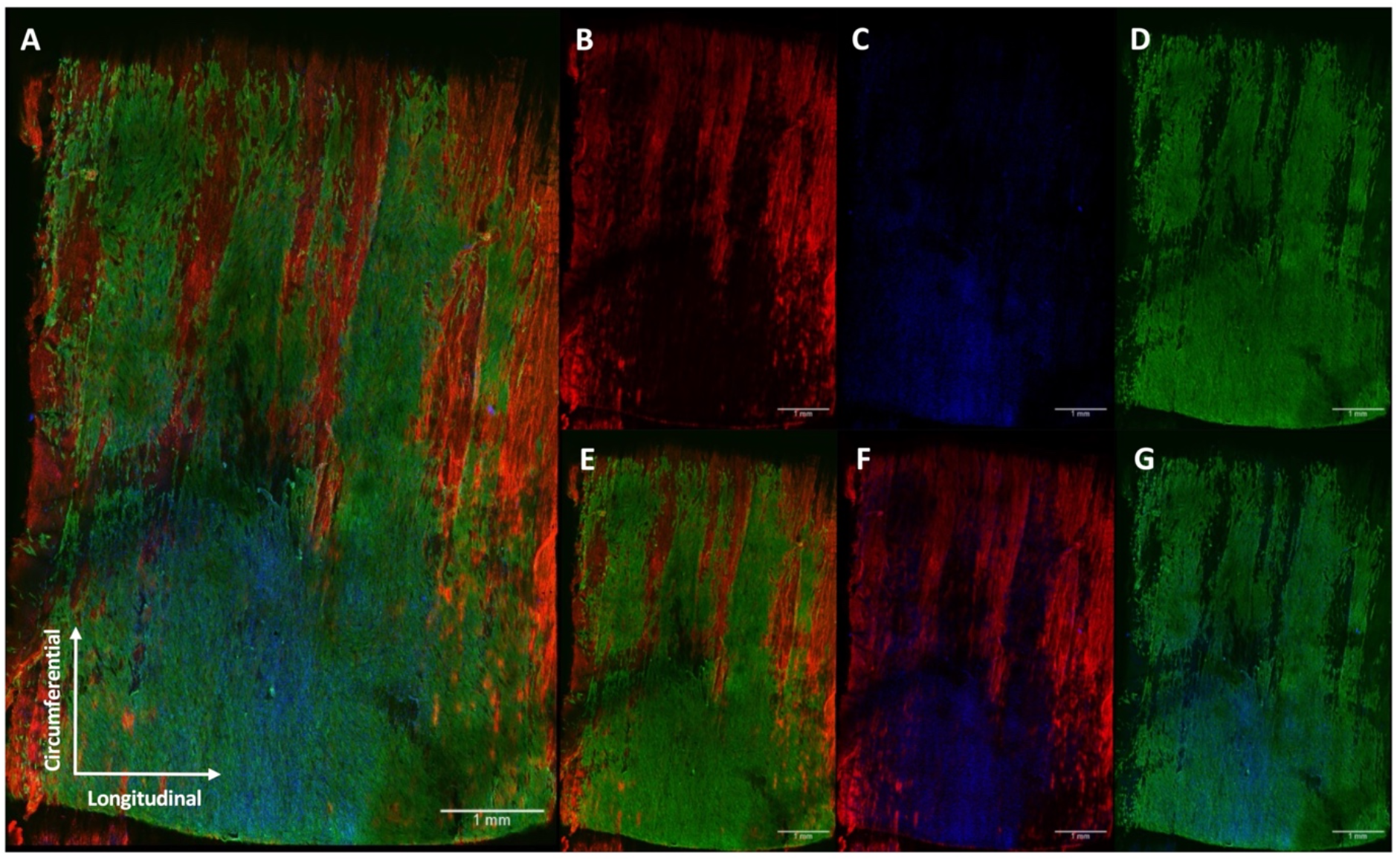
Cell seeding of decellularised vessels. Confocal imaging of pbMSC seeded on decellularised + DNAsed strips stained with CNA35-dTom, SM22α-488 and DAPI following static culture for 48 hrs and 2 – 8% strain for 72 hrs. Scale bars = 1 mm. The vertical axis is the vessels circumferential direction, and the horizontal axis is the longitudinal direction. (A) CNA35-dTom, SM22α-488 and DAPI, (B) CNA35-dTom, (C) DAPI, (D) SM22α-488, (E) CNA35-dTom and SM22α-488, (F) CNA35-dTom and DAPI, (G) SM22α-488 and DAPI.

**Figure 11.**
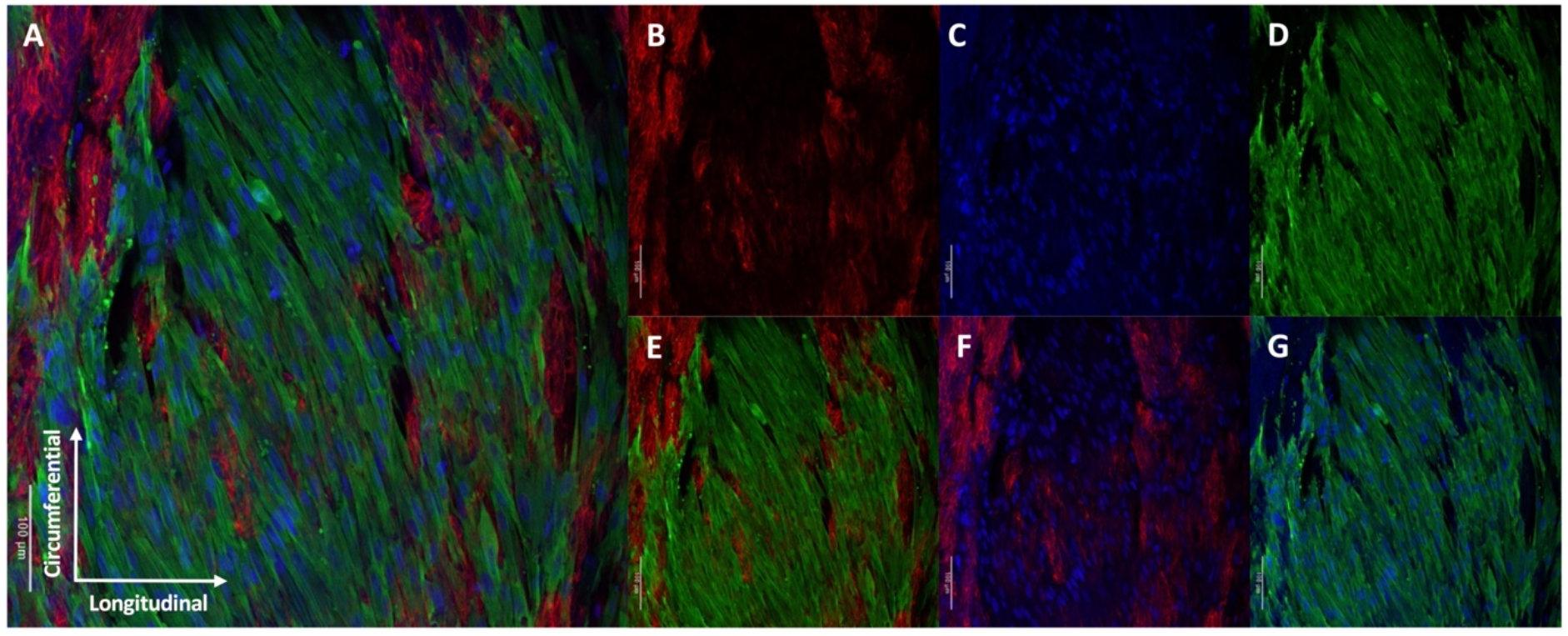
Cell seeding of decellularised vessels. Confocal analysis of pbMSC seeded on decellularised vessels stained with CNA35-dTom, SM22α-488 and DAPI following static culture for 48 hrs and 2 – 8% strain for 72 hrs. Scale bars = 100 *μ*m. The vertical axis is the circumferential direction, and the horizontal axis is the vessels longitudinal direction. (A) CNA35-dTom, SM22α-488 and DAPI, (B) CNA35-dTom, (C) DAPI, (D) SM22α-488, (E) CNA35-dTom and SM22α-488, (F) CNA35-dTom and DAPI, (G) SM22α-488 and DAPI.

**Figure 12.**
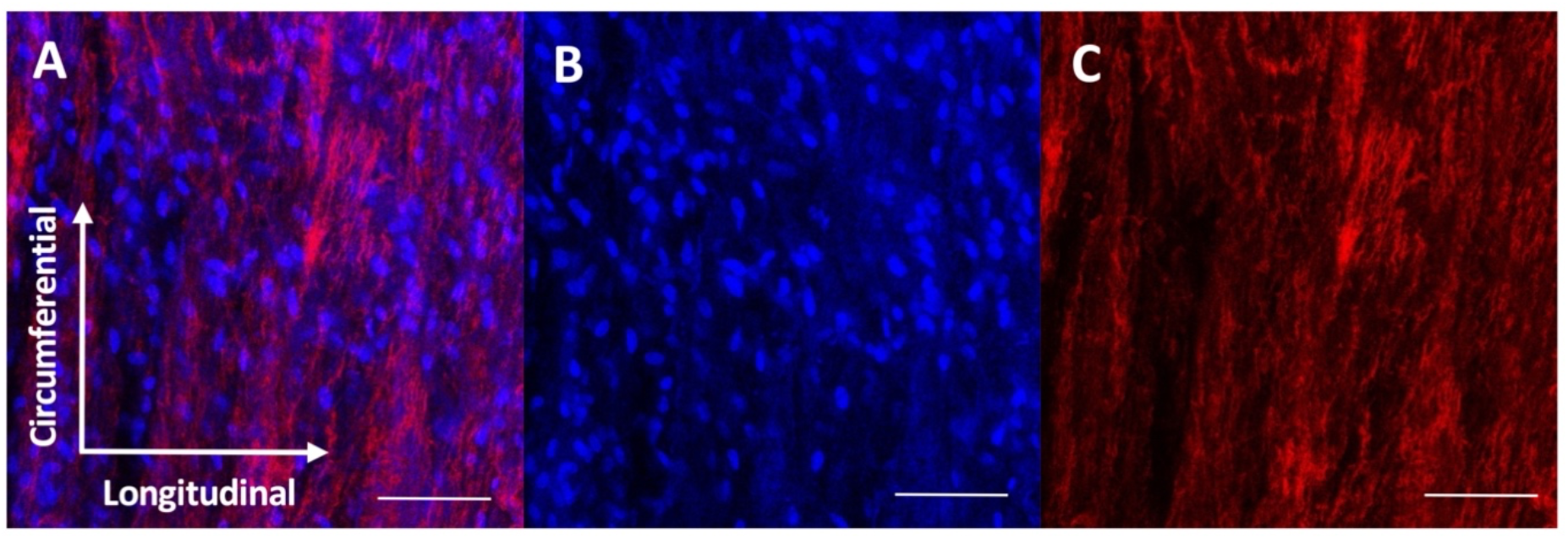
Cell seeding of tubular decellularised vessels. Confocal analysis of pbMSC seeded onto tubular decellularised vessels stained with CNA35-dTom, and DAPI following static culture for 11 days. Scale bars = 100 *μ*m. The vertical axis is the vessels circumferential direction, and the horizontal axis is the vessels longitudinal direction. (A) CNA35-dTom and DAPI, (B) DAPI, (C) CNA35-dTom.

## 4.0 Discussion

Despite significant advances in the field of tissue engineering, the clinical requirement for a biocompatible, patent, structurally and biomechanically vascular mimetic graft still remains considerably unfulfilled [6, 7, 9, 11, 29-33]. While there are currently many successfully developed alternatives showing promise *in vitro*, a reliable alternative to autologous vessels has yet to be found [33-36]. The protocol presented in this paper shows significant promise in addressing this unmet need as it produces grafts which adhere to all the requirements of an “ideal” graft and can be produced in a reliable and efficient manner to yield an off-the-shelf, ready to use product.

Prior to decellularisation, arteries are typically frozen to facilitate long-term storage for convenience purposes. The technique of cryopreservation was originally developed for the long-term storage of isolated cells, and cryopreservation of tissues poses challenges to internal structure, compartmental organisation and cell types [37]. Given the later application of decellularisation, the preserved cell viability aspect of cryopreservation is redundant; however, this study as well as the others suggest that the initial act of preserving cell viability could be key to preventing ECM damage [38, 39]. Like decellularisation, variations in cryopreservation methods are plentiful and can depend on specific characteristics of the tissue being preserved. For vascular tissues, ideal cryopreservation protocols are characterised by the maintenance of vascular functionality post thawing, and comparative mechanical testing of frozen tissues is frequently implemented to determine the effect of freezing on the native mechanical properties [20, 21, 25, 40, 41]. However, depending on the specific vessel characteristics and proposed application after freezing, different preservation conditions are implemented [42, 43]. Cryoprotectant agents are commonly used in conjunction with nourishing vehicle solutions to both support the native cell population and prevent ECM disruption [44]. Other studies have previously reported that unprotected freezing of arteries cause vascular SMCs to show various signs of cell stress, and the absence of a cryoprotectant is known to induce ice crystal formation which in turn disrupts cell membrane integrity and damages surrounding tissue [25, 38]. This study utilised a previously published cryopreservation method specifically for human vessels and confirmed that cryopreservation does not impact vascular structure (Figure 1-Figure 3) [45]. Other studies involving carotid and pulmonary arterial grafts have also shown that cryopreservation preserves tissue structure and results in minimal ECM disruption [39]. Specifically, Chang et al. revealed minimal disruption to the native ECM and cells in carotid arteries and moreover, reported normal nuclear and cytoplasmic morphologies like that observed in this study (Figure 1) [39].

Using an amended version of the decellularisation protocol described previously for coronary arteries by Campbell et al., this study shows comprehensive cellular disruption was achieved across native and cryopreserved sections of porcine carotid arteries after 15 hours of treatment [22]. In comparison, Campbell et al. reported that after 12 hours decellularisation of porcine coronary arteries, all cell nuclei were appropriately disrupted with a total absence of SMCs in the vessel wall [22]. In addition to the larger wall thickness, the increased numbers of elastic lamina and the heterogeneity of the ECM within the medial layer of the elastic carotid artery is likely to have impeded effective infiltration of the decellularisation agent. Nevertheless, in comparison to previous PCA decellularisation protocols, there was no ECM disruption observed in all decellularised vessel states in this study (Figure 1-Figure 3, Figure 5) [24, 46]. Ketchedjian et al. also reported identical tissue structures after decellularisation of both native and cryopreserved pulmonary tissue, a finding that is comparable to this study [47]. However, unlike this study, Ketchedjian et al. did not investigate the impact of the decellularisation technique on the resulting grafts’ biomechanical properties (Figure 8-Figure 9) [47]. Regardless, other studies report imbalances between effective removal of cellular and genetic material, and disruption to ECM structure [24]. One example, published by Sheridan et al., also successfully decellularised frozen porcine carotid arteries using a combination of enzymatic and chemical detergents, but produced a highly porous graft [24]. In that study, H&E staining confirmed an acellular scaffold with high porosity throughout the entire wall thickness [24]. It is also worth noting that the Sheridan et al. protocol took 6 days to produce an acellular construct, whereas the protocol outlined in this study takes significantly less time (2.7 days) to achieve complete removal of cellular content [24]. Additionally, further examination showed no significant difference in collagen fibre orientation, and a single family of predominantly circumferentially oriented collagen fibres was identified in decellularised and DNAse treated vessels (Figure 6) [48]. Critically, collagen content quantification also conducted in this study shows that decellularisation and DNAse treatment does not impact percentage collagen content (Figure 5B).

Following confirmation that the decellularisation protocol outlined in this paper does not impact the underlying collagen structure or content, potential grafts must also comply with decellularisation criteria. Widely accepted decellularisation criteria involve confirming the absence of contaminating cellular and genetic material [18, 19]. H&E examination conducted in this study shows an absence of cellular, but not genetic, material in all decellularised samples (Figure 1). Additionally, examination of DNA levels confirmed the requirement for an additional DNAse treatment procedure. Histological analysis following DNAse treatment qualitatively indicated the absence of DNA following treatment, whilst DNA quantification confirmed sufficient genetic material reduction and to levels within the widely accepted criteria of < 50ng DNA/mg of tissue (dry weight) (Figure 1 and Figure 4) [18, 19]. Additionally, glycosaminoglycans (GAGs) contribute to a variety of functions, including ECM hydration and structure, biochemical signalling, and cell adhesion [58, 59]. The histochemical stain Alcian Blue is a widely used method to visually assess the presence of GAGs and is typically used to qualitatively determine GAG retention following decellularisation [58-62]. Importantly, Alcian Blue staining confirmed that the GAG profile of the vessel remained intact (Figure 3).

Importantly, the international standard for implantable vascular prostheses as described in EN ISO 7198:2017 dictates that water permeability and burst strength characterisation are also required as part of the minimum testing requirement of vascular grafts. The ideal vascular graft must be permeable/porous enough to support molecular transport but have low enough permeability/porosity to prevent haemorrhage and enable anticoagulant administration [27, 49]. The rate of water permeability across the resulting scaffolds from this study (0.009 ± 0.006 ml/cm^2^/min^-1^) complies with this requirement (Figure 7). Unfortunately, there are conflicting reports regarding acceptable permeability/porosity limits, while Wesolowski and McMahon state that the safe limit with regards to haemorrhage is in the vicinity of 5,000 ml/cm^2^/min^-1^, Jonas et al. state that porosity must be controlled to less than 50 ml/cm^2^/min^-1^ to avoid massive haemorrhage [27, 49]. In addition, Madhaven et al. report obtaining a graft with a “desirable” water permeability within the “ideal range of 500-600 ml/cm^2^/min^-1”^ [50]. Considering this, the graft reported in this study is arguably a suitable vascular graft due since it allows water permeation without any danger of haemorrhage. Interestingly, Madhaven et al. also state that water permeability values above 800 ml/cm^2^/min^-1^ require preclotting prior to implantation and below 600 ml/cm^2^/min^-1^ do not [27, 50, 51]. Preclotting is an option to reduce porosity/permeability; however, Jonas et al. warns of a lack of reliability in the technique in certain instances [27, 49, 52]. Nevertheless, the data presented in this paper shows that the grafts resulting from this protocol do not require preclotting and are therefore both convenient and reliable. Additionally, pressurised tests conducted on all scaffolds following the water permeability testing confirmed that there is little or no risk of potential graft integrity failure *in vivo*. Moreover, the grafts exhibited typical vascular non-linear elasticity when mechanically assessed, and the arteries demonstrated the expected exponential strain-stiffening behaviour for increasing circumferential distension (Figure 8). Interestingly, differences were observed between tissue samples obtained from the proximal and distal end of the carotids, which correlates with the structural differences observed during histological analysis (Figure 2 and Figure 8). While it has been previously reported that the collagen:elastin ratio is dependent on arterial type and distance from the heart to the best of the authors knowledge, this observed difference in collagen-elastin ratio has not been previously reported for the porcine carotid artery [53, 54]. The mechanical properties obtained from the uniaxial tensile testing data show that the resulting grafts have preserved mechanical integrity profiles. In addition, the difference in mechanical properties observed, which depends on the vessel location (proximal or distal), could be utilised to better match host vessel mechanical properties, and minimise graft-vessel compliance mismatch (Figure 9).

Irrespective of the scaffolds maintained structural integrity, compliance with decellularisation criteria and appropriate biomechanical profile, if the resulting scaffold cannot host cells, it is of limited value. Therefore, to complete this study, pbMSC were cultured onto both planar strips and tubular constructs of the resulting graft. The tunica media was exposed as it is the most significant load bearing component of the vessel and it provides circumferentially aligned collagen for cell attachment [64, 65]. Unlike other grafts, confocal analysis shows that pbMSC can attach to the graft in either strip or tubular form without the requirement for coating with attachment proteins, which can be both costly and time-consuming (Figure 10-Figure 12) [55-57]. The tubular constructs were cultured for 11 days confirming that the grafts support cell culture over a longer timeframe (Figure 12). Additionally, strips were also subjected to physiological cyclic strain in vitro, and it is noteworthy that the pbMSC maintained attachment despite cyclic strain of 2-8 % being applied over 3 days, confirming that graft repopulation in an in vivo environment is possible (Figure 10-Figure 11).

## Conclusion

In this study, an effective short-term decellularisation protocol was developed for carotid arteries that maintained structural and biomechanical integrity and complied with widely accepted decellularisation criteria. Specifically, the cryopreservation, decellularisation and DNAse treatment did not impact native ECM structure or content and cellular and genetic material could be sufficiently removed. The resulting grafts also demonstrated that they would not haemorrhage or burst under physiological conditions and had appropriate mechanical properties. Finally, pbMSC could be cultured and supported *in vitro* on the resulting graft, therefore demonstrating the potential for graft repopulation under physiological environmental conditions. Critically, this protocol is cost-effective, rapid, automatable and produces a graft that can be used “off the shelf”, as the graft does not require any manipulation to promote cell attachment or pre-clotting to prevent haemorrhaging. Ultimately, it is proposed that the resultant acellular tubular scaffold presents a great potential alternative to autologous grafts.

## Acknowledgements

This publication has emanated from research conducted with the financial support of the European Research Council (ERC) under the European Union’s Horizon 2020 research and innovation programme (grant agreement no. 637674) and Science Foundation Ireland under the grant no. SFI/13/ERC/B2775.

