## Supplementary figures 1-3 for "A short-term decellularisation technique for porcine carotid arteries that conveys a structural stimulus to cells"

[1]Trinity Centre for Biomedical Engineering, Trinity Biomedical Sciences Institute, Trinity College Dublin, Dublin 2, Ireland; [2]Department of Mechanical, Manufacturing & Biomedical Engineering, School of Engineering, Trinity College Dublin, Dublin 2, Ireland; [3]School of Biotechnology, Vascular Biology & Therapeutics Group, Dublin City University, Glasnevin, Dublin 9, Ireland; [4]Advanced Materials and Bioengineering Research Centre (AMBER), Royal College of Surgeons in Ireland and Trinity College Dublin, Dublin 2, Ireland.

### Supplementary material

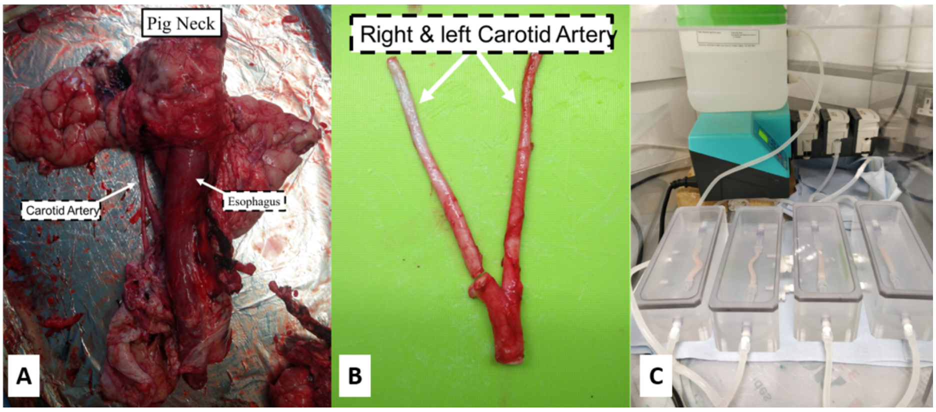


**Supplementary figure 1:** Porcine carotid artery dissection and decellularisation. (A) Location of carotid artery in a pig neck, (B) right and left carotid post dissection, and (C) the decellularisation rig.

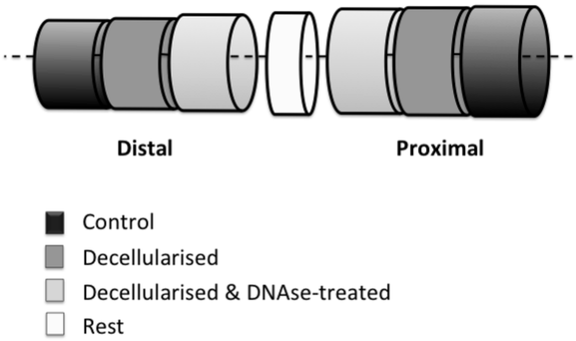


**Supplementary figure 2:** Diagram of sample location in mechanical testing experiments. Proximal and distal sample locations were taken for each treatment.

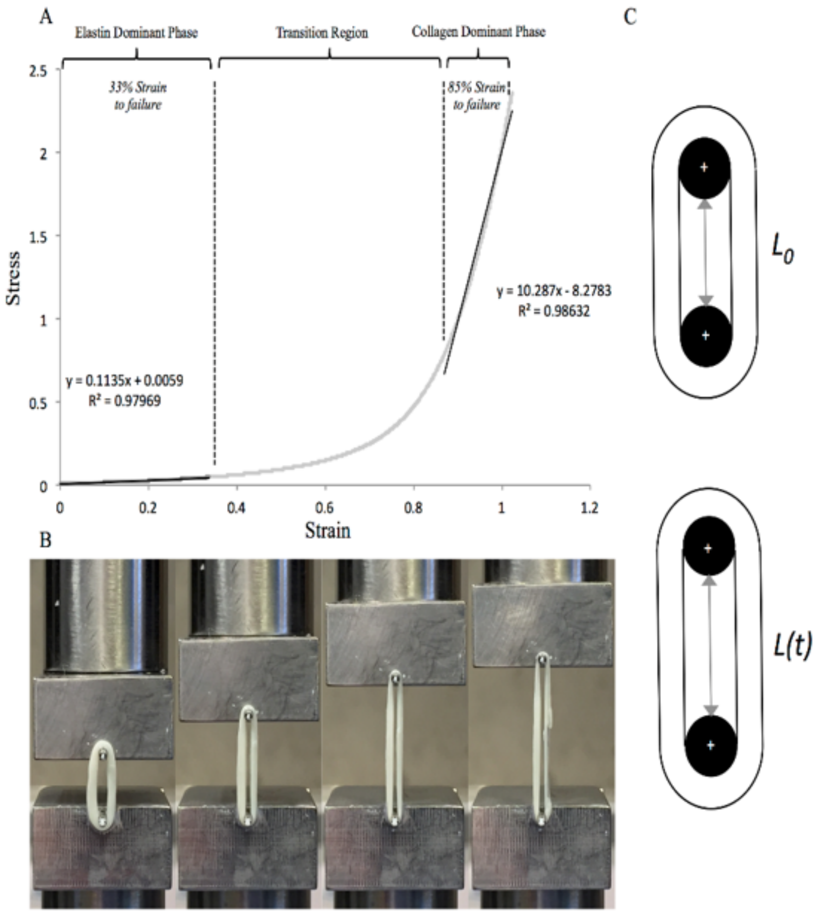


**Supplementary figure 3:** Representative stress-strain plot and corresponding sample elongation. (A) Example stress-strain plot for uniaxial tensile test of PCA ring sample, (B) Macroscopic progression of sample elongation after preconditioning (LHS) up to failure (RHS), (C) Schematic showing initial and gauge length at time t.
